# Highly diverse fungal communities in carbon-rich aquifers of two contrasting lakes in Northeast Germany

**DOI:** 10.1101/623272

**Authors:** Anita Perkins, Lars Ganzert, Keilor Rojas-Jiménez, Jeremy Fonvielle, Grant C. Hose, Hans-Peter Grossart

## Abstract

Fungi are an important component of microbial communities and are well known for their ability to degrade refractory, highly polymeric organic matter. In soils and aquatic systems, fungi play an important role in carbon processing, however, their diversity, community structure and function as well as ecological role, particularly in groundwater, are poorly studied. The aim of this study was to examine the fungal community composition, diversity and function of 16 groundwater boreholes located in the vicinity of two lakes in NE Germany that are characterized by contrasting trophic status. The analysis of 28S rRNA gene sequences amplified from the groundwater revealed high fungal diversity and clear differences in community structure between both aquifers. Most sequences were assigned to *Ascomycota* and *Basidiomycota*, but members of *Chytridiomycota, Cryptomycota, Zygomycota, Blastocladiomycota, Glomeromycota* and *Neocallmastigomycota* were also detected. In addition, 27 species of fungi were successfully isolated from the groundwater wells and tested for their ability to degrade complex organic polymers – the predominant carbon source in the wells. Most isolates showed positive activities for at least one of the tested polymer types, with three strains, belonging to the genera *Gibberella, Isaria* and *Cadophora*, being able to degrade all tested substrates. Our results highlight the high diversity of fungi in groundwater, and point to their important ecological role in breaking down highly polymeric organic matter in these isolated microbial habitats.

## Introduction

Aquifers are a key component of Earth’s hydrological cycle and form the largest accessible freshwater reservoir on the planet (Danielopol et al., 2003). Through the flow of groundwater, aquifers constitute an important pathway between terrestrial and aquatic ecosystems and can be a sink or a source of dissolved organic matter (DOM) for these ecosystems.

Organic matter (OM) from terrestrial systems can be dissolved by rainfall and percolate through the soil profile and into the groundwater (Chapelle, 2001). The majority of the DOM in surface waters is made up of polysaccharides, lignin, proteins, and lipid sugars that originate mainly from phytoplankton (Grossart and Simon, 2007) and plant material (Bertilsson and Tranvik, 2000; Rochelle-Newall and Fisher, 2002; Zhang et al., 2007), but DOM can also be produced or modified *in-situ* (Shen et al., 2015; Rojas-Jiménez et al., 2017). The concentration and chemical structure of OM in groundwater systems depends mainly on the geological complexity and sorption characteristics of the sediment layers, the type of aboveground vegetation, land use and microbial activity in the soil and aquifer sediments (Jardine et al., 1989; Williams et al., 2010; Peter et al., 2012; Chapelle et al., 2013; Graham et al., 2015). These factors typically influence the concentrations of coloured DOM (CDOM) and fluorescent DOM (FDOM) in the overall DOM pool, which in turn determines the overall OM quality in groundwater (Mladenov et al., 2010; Li et al., 2014). Although DOM in groundwater can be a major carbon source for surface water systems (Cai et al., 2003; Mitch and Gosselink, 2007; Santos et al., 2012; Maher et al., 2013), aquifers are rarely considered as part of the global C cycle (Macpherson, 2009).

In soils, fungi are important OM decomposers, particularly of biological polymers such as lignin, cellulose and chitin, and as such contribute significantly to global biogeochemical cycles (Dashtban et al. 2010; Treseder and Lennon 2015). While fungi and bacteria often compete for the same resources, the physiological capabilities of fungi makes them more effective in breaking down refractory OM (Jørgensen and Stepanauskas, 2009; Harms et al., 2011; Rojas-Jiménez et al., 2017). Fungi are also an ecologically important and diverse component of groundwater microbial communities (Lategan et al., 2012; Lategan and Hose, 2014; Sohlberg et al., 2015; Nawaz et al., 2016, 2018; Korbel et al., 2017). Moreover, groundwater can be a conduit for the exchange of spores and propagules between surface waters and soils (Brad et al., 2008).

Despite their importance to global geochemical cycles, aquatic fungi remain a poorly studied group with thousands of species yet to be identified (Gessner, 1997; Wurzbacher et al., 2010; Grossart et al., 2015; Grossart and Rojas-Jiménez, 2016). In aquifers, even less is known about the diversity and functions of fungi, and their interactions with other organisms. Methodological difficulties involved with culturing and the identification of mycelial growth forms, unclear taxonomy and limited databases make fungi difficult to profile accurately in many environments. In addition, the processes by which fungi use organic matter in aquatic habitats and their role in ecosystem balance remains poorly understood (Clipson and Gleeson, 2012; Crowther and Grossart, 2015), especially in groundwater environments (Sohlberg et al., 2015; Nawaz et al., 2016, 2018).

In this study, we investigated the diversity and structure of fungal communities in groundwater in the vicinity of two lakes in Northeast Germany that differed markedly in their trophic status. We hypothesized that, given the differences in water quality between the aquifers surrounding the lakes, fungal communities would also differ. We further investigated the potential of isolated groundwater fungal taxa to degrade highly polymeric and potentially fossil carbon sources, expecting that different taxa would differ in their degradation ability. We found a high fungal diversity in both aquifers, and clear differences in community structure between the aquifers. Many of the fungal isolates showed positive activities for at least one of the tested polymer types. These results highlight that aquifers are a suitable fungal habitat and indicate an important role of groundwater fungi in the breakdown of highly polymeric OM.

## Material and Methods

### Study Site

Groundwater samples were collected from 11 boreholes around Lake Stechlin and 5 boreholes around Lake Grosse Fuchskuhle (Figure 1), located in northeast Germany. Both lakes and catchments are surrounded by forest and their watersheds are relatively little impacted by human activities. Lake Stechlin is a dimictic calcareous lake covering 4.25 km^2^, with a maximum depth of 69.5 m, a volume of 96.9 GL and a catchment area of 42.3 km^2^ (Casper, 1985). Lake Grosse Fuchskuhle is a naturally acidic bog lake that covers ca. 0.02 km^2^, with a maximum depth of 5.6 m, a volume of 0.05 GL and a catchment area of 0.005 km^2^ (Burkert et al., 2004; Allgaier and Grossart, 2006).

**Fig 1:**
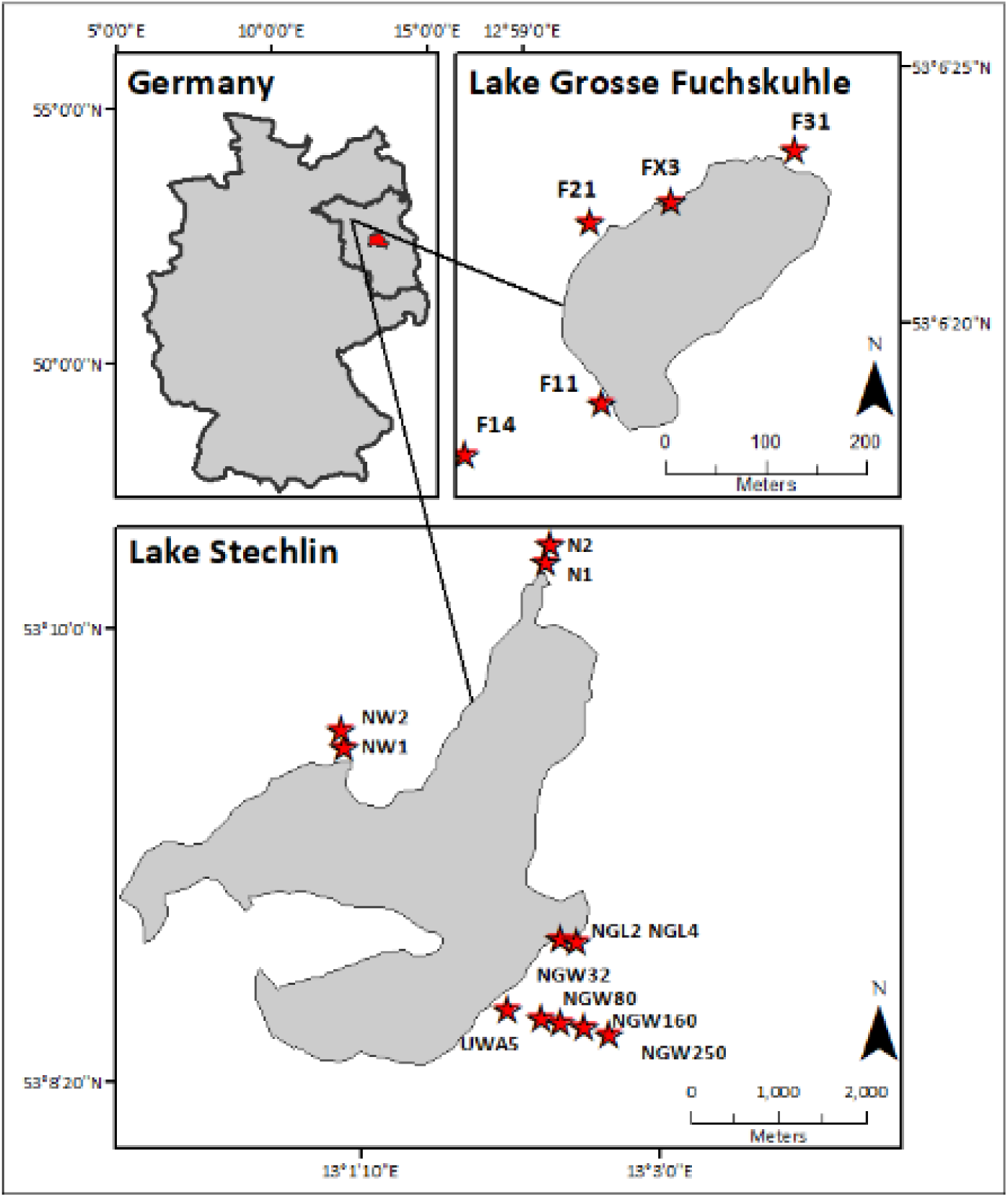
Location of the boreholes surrounding Lake Stechlin and Lake Grosse Fuchskuhle, ca. 80 km North of Berlin (Germany).

The two lakes differ greatly in their trophic status: Lake Stechlin is oligo-/mesotrophic (pH 7.2 - 8.5), while Lake Grosse Fuchskuhle is a dystrophic lake with a pH of < 6 (Allgaier and Grossart, 2006), receiving large amounts of humic substance (HS) from a surrounding bog system (Hutalle-Schmelzer et al., 2010). Both, Lake Stechlin and Lake Grosse Fuchskuhle were formed by glacial dead ice blocks and have no contribution from rivers or streams, thus the lake’s water is derived solely from groundwater and precipitation (Ginzel, 1999; Holzbecher et al., 1999). Further details can be found in the supplementary information.

### Sampling procedure

Water samples were collected in May 2016. Boreholes were carefully sampled using a surface-sterilized submersible pump (MP1, Grundfos) and were purged prior to sample collection by removing at least twice the total borehole volumes.

Water samples for chemical analysis were collected using a polypropylene syringe (60 mL, pre-rinsed with borehole water) and stored in acid-washed 20 mL polyethylene vials. Water samples were analysed for dissolved organic carbon (DOC), dissolved inorganic carbon (DIC), soluble reactive phosphorus (SRP), nitrite (NO_2_), nitrate (NO_3_), ammonium (NH_4_), iron (Fe), total nitrogen (TN) and total phosphorus (TP). Samples for TN and TP analysis were analysed unfiltered. Samples for all other analyses were filtered using a 0.45 µm pore-size syringe filter (pre-rinsed with 60 mL Milli-Q water). Samples for ^18^O and Deuterium (^2^H) measurements were collected into 2 mL sterile glass vials. Temperature, pH, electrical conductivity and dissolved oxygen of the groundwater were recorded on site using a YSI probe (Yellow Springs Instruments).

Water samples for molecular analysis were collected in sterile 2.5 L HDPE bottles and kept dark at 4°C in a cooler. Samples were filtered in the laboratory within <4 h of collection using 0.2 µm pore-size Sterivex filters. Up to 1.5 L of sample was filtered until the filter became clogged. Filter membranes were stored at −20°C until further analysis.

### Chemical analysis

DOC concentrations were determined using a TOC-Vcph total organic carbon analyser (Shimadzu). DIC was determined as total dissolved carbon minus DOC, which has been determined after acidification to remove all DIC before the measurement. SRP, TP, TN, NO_2_, NO_3_, NH_4_ and Fe were measured photometrically using a FIASTAR 5000 (Foss Analytical AB).

### Optical characterisation of dissolved organic matter (DOM)

DOM was characterized optically using a UV-Vis spectrophotometer (Hitachi U-2900) to measure absorbance between 190 and 800 nm in 1 nm steps. If the absorbance was higher than 0.3 at 300 nm, the samples were diluted with Milli-Q water to reduce the inner filter effect. Excitation-emission matrices (EEM) were obtained using a fluorescence spectrophotometer (Hitachi F-7000) with excitation ranging from 220 to 450 nm (5 nm increment) and emission ranging from 230 to 600 nm (2 nm increment). EEMs were corrected against a Milli-Q water sample and further interpolations of primary and secondary scatters of Rayleigh and Raman regions were performed prior to correction of the remaining inner filter effect using an absorbance-based method (Christmann et al., 1980; Murphy et al., 2013). Further details of defining DOM fractionation, characteristics and calculations are provided in the supplementary information.

### DNA extraction and fungal diversity assessment by amplicon sequencing

Genomic DNA from Sterivex filters (Millipore Corporation) was extracted using a CTAB-phenol-chloroform-isoamylalcohol/bead beating protocol (modified after Nercessian et al., 2005). Details of the extraction procedure can be found in the supplementary information. PCR, library preparation and sequencing was performed by LGC Genomics (Berlin, Germany). Briefly, the D2 region was amplified using primers LR22R-LR3 (Mueller et al., 2016), followed by library preparation (2×300 bp) and sequencing on a MiSeq Illumina platform. Sequences were quality checked and analysed using Mothur v1.37.6 (Schloss et al., 2009). Sequences shorter than 150 bp or which contained ambiguities and homopolymer stretches of more than 8 bases were excluded from further analyses. A chimera check was performed using UCHIME (Edgar et al., 2011) in *de-novo* mode. Sequences were clustered into operational taxonomic units (OTU) using VSEARCH (Rognes et al., 2016; as implemented in Mothur) with a minimum sequence similarity value of 97%. Global singleton sequences were removed. Taxonomy assignment of the OTUs was based on Fasta36 (Pearson, 2002), using the SILVA LSU reference database v128. The five best hits to the highest common taxonomy level were parsed using a custom-made Python script. All non-fungal taxa were removed from the OTU table. For alpha diversity calculation all samples were subsampled to 500 sequence reads (sample NGL4 was removed due to too few useful sequence reads). All sequence reads are available in the Sequence Read Archive (SRA) under the BioProject PRJNA485560.

### Fungal isolation and molecular identification

Groundwater (250 mL) from three selected boreholes (NW1, NGW250, FX3) was filtered through 5.0 µm polycarbonate membranes (Millipore, USA) using aseptic conditions. The filter contents were re-suspended in 1 mL PBS 1X (10 mM Na_2_HPO_4_, 1.8 mM KH_2_PO_4_, 137 mM NaCl, and 2.7 mM KCl) and plated on cultivation media (Rojas-Jiménez et al., 2017) amended with each 100 mg L^−1^ of streptomycin and penicillin. Plates were incubated at room temperature (20°C) for up to seven days. Emerging fungi were separated and cultured on new plates. After two weeks of growth, we compared the morphotypes of each isolate, considering characteristics such as color, texture, shape, and growth. For all isolates showing identical characteristics duplicate cultures were removed.

DNA was extracted from ca. 250 mg of fungal mycelia using the peqGOLD Tissue DNA Mini Kit (Peqlab, Germany). The ITS1-5.8S-ITS2 region was amplified with primers ITS1 and ITS4 (White et al., 1990), and a fragment of the 28S rRNA gene using primers LROR-LR5 (Vilgalys and Hester, 1990). PCR was performed in a 50 µL reaction, using MyTaq Red DNA Polymerase (Bioline, Germany) with the following reaction conditions: 94°C for 2 min, 32 cycles at 94°C for 15 sec, 53°C for 15 sec, 72°C for 30 sec, and a final extension at 72°C for 5 min. PCR products were Sanger sequenced at Macrogen Europe. Sequences were assembled using BioEdit (Hall, 1999).

Fungal identification was performed using different reference databases for each marker: for the ITS region we used GenBank, the Warcup Fungal ITS training set 2, and the UNITE Fungal ITS training set. A maximum likelihood phylogenetic tree of the concatenated alignments of LSU+ITS was generated with FastTree version 2.1.9 (Price et al., 2010), using a GTR+G+I model of evolution. The resulting phylogenetic tree was displayed and edited in MEGA7 (Kumar et al., 2016). Further details are provided in the supplementary information. Sequences are available in GenBank under the accession numbers MK007260-MK007286 (LSU) and MK012396-MK012422 (ITS).

### Polymeric substrate degradation by fungal isolates

The ability to degrade polymeric substrates was tested in laboratory assays. Assays were performed by inoculating each fungal strain on 6-well plates with cultivation medium amended with one of the following substrates: 1) 0.1% wt/vol ABTS (2,2′-Azino-bis 3-ethylbenzothiazoline-6-sulfonic acid di-ammonium salt), 2) 0.02% wt/vol Remazol Brilliant Blue (RBBR), 3) 0.02% wt/vol Bromocresol Green (Bromo), 4) 0.02% wt/vol PolyR-478 (PolyR), 5) 0.02% wt/vol Toluidine Blue (Tol), and 6) 0.02% wt/vol Congo Red (Congo). Capability of degrading ABTS was detected as a color change demonstrating laccase activity. Decolorization of Remazol Brilliant Blue R (RBBR) is related to lignin peroxidase activity, while decolorization of PolyR-478 (PolyR), Bromocresol Green (Bromo), Toluidine (Tol), and Congo Red (Congo) is related to degradation of polymeric, triarylmethane, and heterocyclic substrates, respectively. The capacity for hydrolyzing the substrates was detected as a color change of the media (substrate 1) or as decolorization of the area around the mycelia (substrates 2 to 6) after 3 weeks.

### Statistical analyses

Data processing, visualization and statistical analysis were performed in R (R-Core-Team, 2017). The Vegan package (Oksanen et al., 2017) was used for non-metric multidimensional scaling (NMDS), permutational analysis of variance (PERMANOVA) and Analysis of Multivariate Homogeneity of group dispersions to estimate differences in the fungal community structure using normalized data by converting OTU counts (without singletons) into relative abundances. In order to limit biases due to low sequencing depth, we removed samples with low sequence numbers (N1, NW1, NGL4).

Differences in DOM composition were displayed using a principal component analysis (PCA) after z-transformation of the data. Differences in individual parameters between sites were tested using a Wilcoxon rank test. To visualize OTUs that were unique to Lake Stechlin or Lake Grosse Fuchskuhle or were shared by both lakes, we converted the OTU matrix into presence/absence data. An undirected network was computed using the package ‘igraph’ (Csardi and Nepusz, 2006; http://igraph.org) and visualized with Cytoscape (Shannon et al., 2003). Species richness and evenness were tested for differences by performing Analysis of Variance (ANOVA) and Tukey’s test for pairwise comparisons.

## Results

### Groundwater characteristics

Groundwater characteristics of the boreholes around the two lakes are shown in Table 1. Regardless of lake pH, the pH of the groundwater was always above 6, ranging from pH 6.03 to 7.57. Conductivity ranged from 63 to 472 µS cm^−1^ for Lake Grosse Fuchskuhle and from 284 to 836 µS cm^−1^ for Lake Stechlin, with the exception of sample NGW250 that had a much higher conductivity of 2991 µS cm^−1^ compared to all other samples. Isotope concentrations at Lake Grosse Fuchskuhle ranged from −4.8 to −6.7 [‰]for ^18^O and from −40.53 to 56.6 [‰] for ^2^H whereas at Lake Stechlin the concentrations ranged from −7.7 to −9.2 [‰]for ^18^O and from −55.4 to −63.7 [‰] for ^2^H, with the exceptions of NW1 where ^18^O was −2.9 [‰] and ^2^H −29.7 [‰] and N1 where ^18^O was −2.0 [‰] and ^2^H was −24.3 [‰]. Nitrite and nitrate were not detectable for samples close to Lake Grosse Fuchskuhle. For samples close to Lake Stechlin, nitrite and nitrate concentrations were generally low (≤ 0.045 mg L^−1^ and ≤ 0.1 mg L^−1^, respectively), except for samples NW2, NGL2 and NGL4 were nitrate was relatively high, ranging from 0.7 to 3.6 mg L^−1^.

**Table 1:** Environmental parameters of the groundwater samples. Temp – temperature, Cond – conductivity, DO sat – dissolved oxygen saturation, DOC – dissolved organic carbon, CDOM – coloured dissolved organic matter, DIC – dissolved inorganic carbon, SRP – soluble reactive phosphorous, TP – total phosphorous, TN – total nitrogen, N/A – not available.

| Sample ID | Depth | Temp | Cond | pH | DO sat | DOC | CDOM | DIC | SRP | TP | NO<sub>2</sub> | NO<sub>3</sub> | NH<sub>4</sub> | TN | Fe | <sup>18</sup>O | <sup>2</sup>H |
| --- | --- | --- | --- | --- | --- | --- | --- | --- | --- | --- | --- | --- | --- | --- | --- | --- | --- |
|  | m | °C | µS cm<sup>-1</sup> |  | mg L<sup>-1</sup> | mg L<sup>-1</sup> | m<sup>-1</sup> | mg L<sup>-1</sup> | mg L<sup>-1</sup> | mg L<sup>-1</sup> | mg L<sup>-1</sup> | mg L<sup>-1</sup> | mg L<sup>-1</sup> | mg L<sup>-1</sup> | mg L<sup>-1</sup> | [‰] | [‰] |
| N1 | 4.9 | 9.3 | 284 | 6.58 | 0.53 | 11.62 | 31.3 | 27.36 | 0.028 | 0.035 | 0.002 | 0.001 | 0.309 | 0.891 | 0.02 | -2.0 | -24.3 |
| N2 | 8.4 | 9.9 | 405 | 6.54 | 0.40 | 18.41 | N/A | 37.63 | 0.067 | 0.105 | 0.001 | 0.003 | 1.522 | 1.938 | 0.517 | -7.7 | -55.4 |
| NGL2 | 1.7 | 9.0 | 771 | 7.25 | 1.43 | 13.75 | 25.3 | 60.67 | 0.004 | 0.015 | 0.043 | 3.647 | 0.037 | 5.432 | 0.005 | -8.9 | -62.8 |
| NGL4 | 4.2 | 8.6 | 836 | 7.25 | 0.81 | 16.06 | N/A | 60.53 | 0.007 | 0.012 | 0.045 | 1.746 | 0.032 | 3.287 | 0.005 | -8.8 | -62.0 |
| NW1 | 4.7 | 9.5 | 458 | 7.40 | 1.26 | 13.23 | 30.1 | 40.90 | 0.006 | 0.016 | 0.001 | 0.002 | 0.139 | 0.441 | 0.098 | -2.9 | -29.7 |
| NW2 | 6.6 | 9.5 | 580 | 7.45 | 7.35 | 12.09 | 4.9 | 49.69 | 0.004 | 0.014 | 0.001 | 0.709 | 0.039 | 1.311 | 0.002 | -8.8 | -61.3 |
| UBA5 | 5.1 | 9.8 | 520 | 7.57 | 1.25 | 16.58 | 29.8 | 50.07 | 0.036 | 0.045 | 0.016 | 0.080 | 0.031 | 0.308 | 0.021 | -8.8 | -61.5 |
| NGW32 | 32 | 9.1 | 517 | 7.49 | 0.73 | 9.31 | 9.5 | 43.95 | 0.008 | 0.014 | 0.001 | 0.101 | 0.047 | 0.247 | 0.004 | -9.2 | -63.7 |
| NGW80 | 87 | 10.8 | 354 | 7.55 | 0.62 | 10.97 | 32.9 | 34.73 | 0.041 | 0.058 | 0.002 | 0.001 | 0.074 | 0.127 | 0.294 | -9.1 | -64.0 |
| NGW160 | 160 | 12.7 | 349 | 7.41 | 1.16 | 14.05 | 34.3 | 40.97 | 0.075 | 0.092 | 0.001 | 0.001 | 0.568 | 0.739 | 0.390 | -8.8 | -61.2 |
| NGW250 | 250 | 14.1 | 2991 | 8.08 | 0.06 | 336.85 | 5437.4 | 78.63 | 0.766 | 0.888 | 0 | 0 | 0.921 | 5.136 | 3.010 | -7.7 | -58.6 |
| F1.1 | 6.5 | 11.2 | 472 | 7.33 | N/A | 28.63 | 171.1 | 58.56 | 0.134 | 0.350 | 0 | 0 | 3.207 | 4.144 | 0.311 | -4.8 | -42.8 |
| F1.4 | 6.0 | 10.4 | 460 | 7.07 | N/A | 81.23 | 727.4 | 64.74 | 0.097 | 0.190 | 0 | 0 | 2.366 | 4.254 | 0.910 | -4.9 | -40.3 |
| **F2.1** | 5.7 | 11.0 | 167 | 6.03 | N/A | 48.77 | 202.7 | 35.19 | 0.053 | 0.082 | 0 | 0 | 2.965 | 4.596 | 1.890 | -8.1 | -56.6 |
| **F3.1** | 6.2 | 10.8 | 419 | 7.21 | N/A | 25.90 | 203.4 | 47.47 | 0.151 | 0.208 | 0 | 0 | 1.408 | 2.329 | 1.764 | -6.7 | -51.0 |
| **FX3** | 4.5 | 13.9 | 63 | 6.25 | N/A | 15.89 | 390.2 | 7.15 | 0.014 | 0.018 | 0 | 0 | 0.497 | 1.114 | 3.729 | -6.4 | -49.1 |

All groundwater samples from Lake Grosse Fuchskuhle had a brownish color, whereas groundwater samples from Lake Stechlin were generally colorless with two exceptions: sample N2 showed a slight yellowish coloration while sample NGW250 was black (see Fig. S1). DOC concentrations of the groundwater differed between the two lakes, with concentrations ranging from 9.31 to 336.85 mg L^−1^ and from 15.89 to 81.23 mg L^−1^ for Lake Stechlin and Lake Grosse Fuchskuhle, respectively (Table 1). The exceptionally high DOC concentration of 336.85 mg L^−1^ for borehole NGW250 was indicated by the black color of the sample; other groundwater samples from around Lake Stechlin were all below 18.5 mg DOC L^−1^.

Although showing a trend for different DOM sources, no significant differences in DOM quality (for HIX, FIX, SR, E2/E3) in the groundwater between Lake Stechlin and Lake Grosse Fuchskuhle were detected (Fig. 2a; Wilcoxon test, p>0.05). The exception was for SUVA_254_ values, which were significantly higher in samples from Lake Grosse Fuchskuhle (4.53 ± 3.58) compared to those from Lake Stechlin (1.62 ± 2.11) (Fig. 2b; Wilcoxon test, p=0.01).

**Fig 2:**
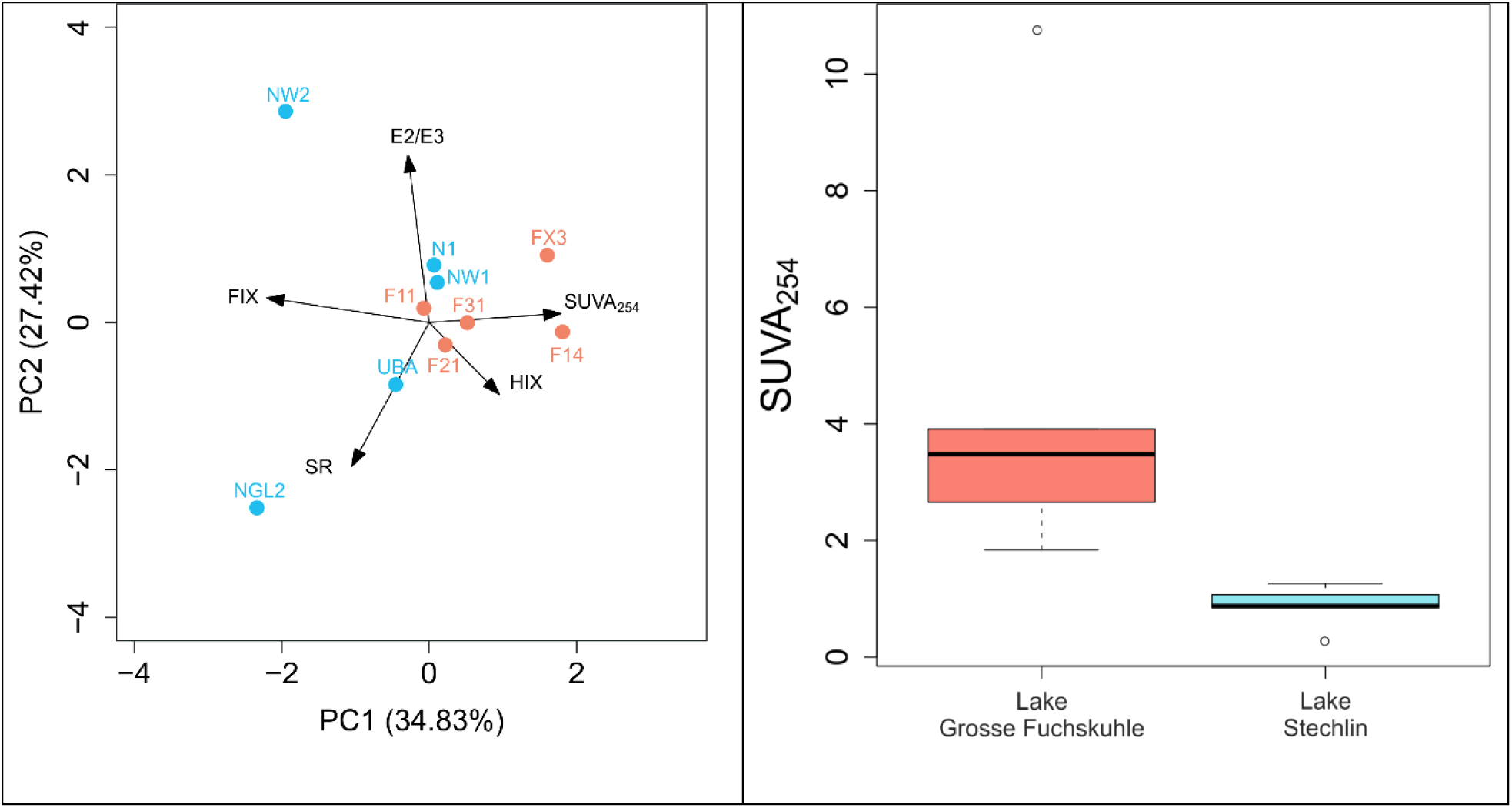
(a) Principal component analysis (PCA) of the shallow groundwater sites based on parameters of DOM quality. The percentage of explained variance of each principal component (PC) is given in brackets. (b) Significant differences in SUVA_254_ values for the same groundwater sites (Wilcoxon test, p=0.01). Box plots show the median value (center line), the upper/lower quartiles (box edges) and maxima/minima (whiskers), and outliers (individual data points). Both sites correspond to n=5. Red - Lake Grosse Fuchskuhle, blue - Lake Stechlin.

### Fungal community structure and diversity

After quality checking, we identified 1070 fungal OTUs out of 143620 sequences. Most sequences were assigned to *Ascomycota* and *Basidiomycota*, but also OTUs belonging to *Chytridiomycota, Cryptomycota, Zygomycota, Blastocladiomycota, Neocallimastigomycota* and *Glomeromycota* were detected (Fig. 3). Sequences belonging to *Cryptomycota* were found in in several samples and reached up to 72% of all fungi in sample FX3. The overall most abundant OTU belonged to the genus *Exobasidium* (*Basidiomycota*).

**Fig 3:**
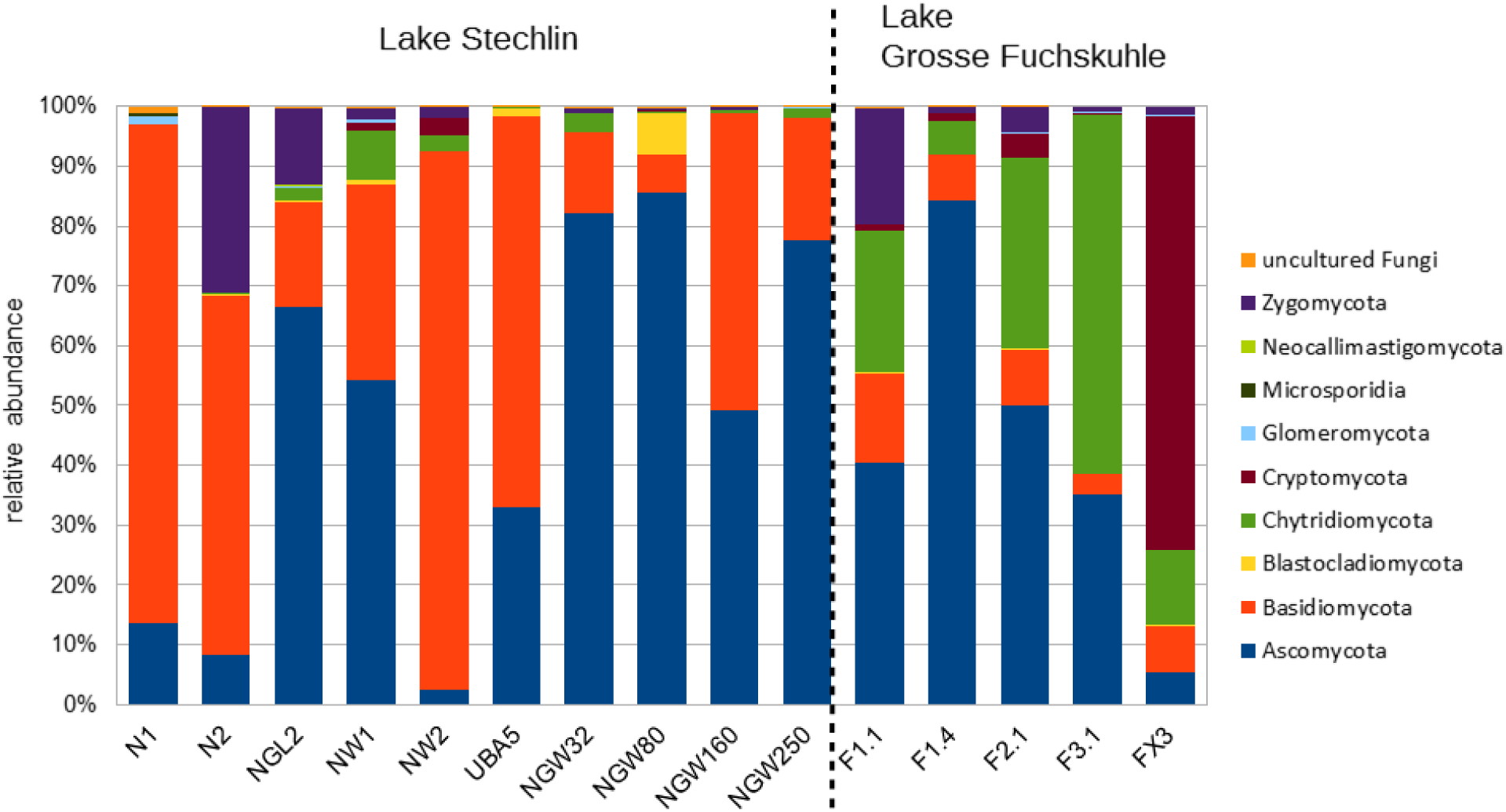
Relative abundance of fungal phyla for the different investigated groundwater samples from the two contrasting lakes. Note: groundwater sample NGL4 from Lake Stechlin is not shown because of too low obtained sequence numbers.

NMDS analysis showed a clear separation of the fungal communities in the aquifers surrounding the two different lakes (Fig. 4), which was confirmed by a PERMANOVA analysis (p=0.002). A similar result was obtained by a network analysis based on presence/absence of fungal OTUs revealing a clear distinction between the groundwater surrounding the two lakes. We found 461 OTUs only in boreholes in the vicinity of Lake Stechlin, 367 OTUs around Lake Grosse Fuchskuhle, while 242 OTUs were common in the groundwater of both lakes (Fig. 5).

**Fig 4:**
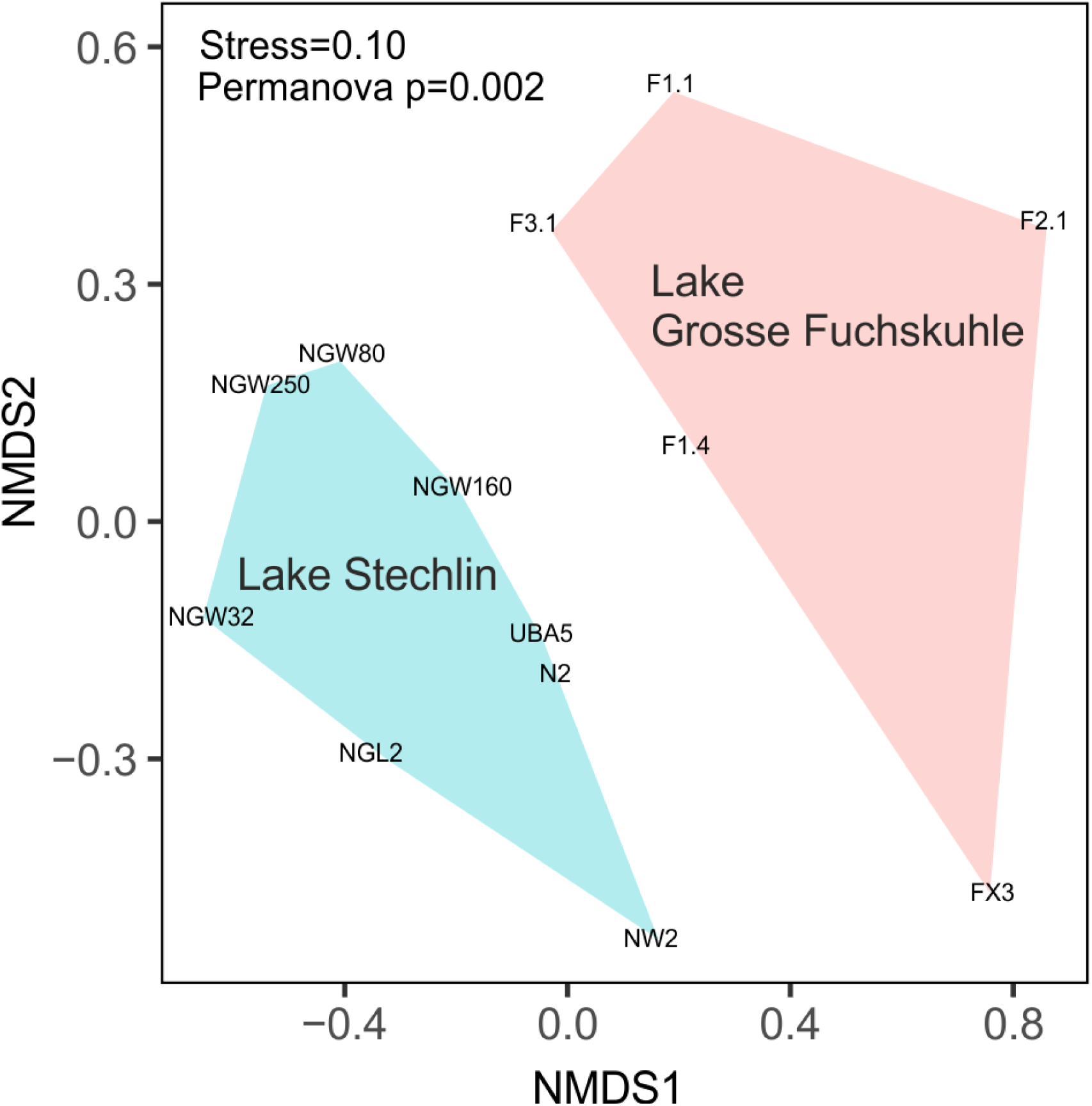
NMDS plot showing the separation of the groundwater fungal community in the vicinity of Lake Stechlin and Lake Grosse Fuchskuhle (based on relative OTU abundances). Polygons refer to groundwater samples from Lake Stechlin (blue) and Lake Grosse Fuchskuhle (red), respectively.

**Fig 5:**
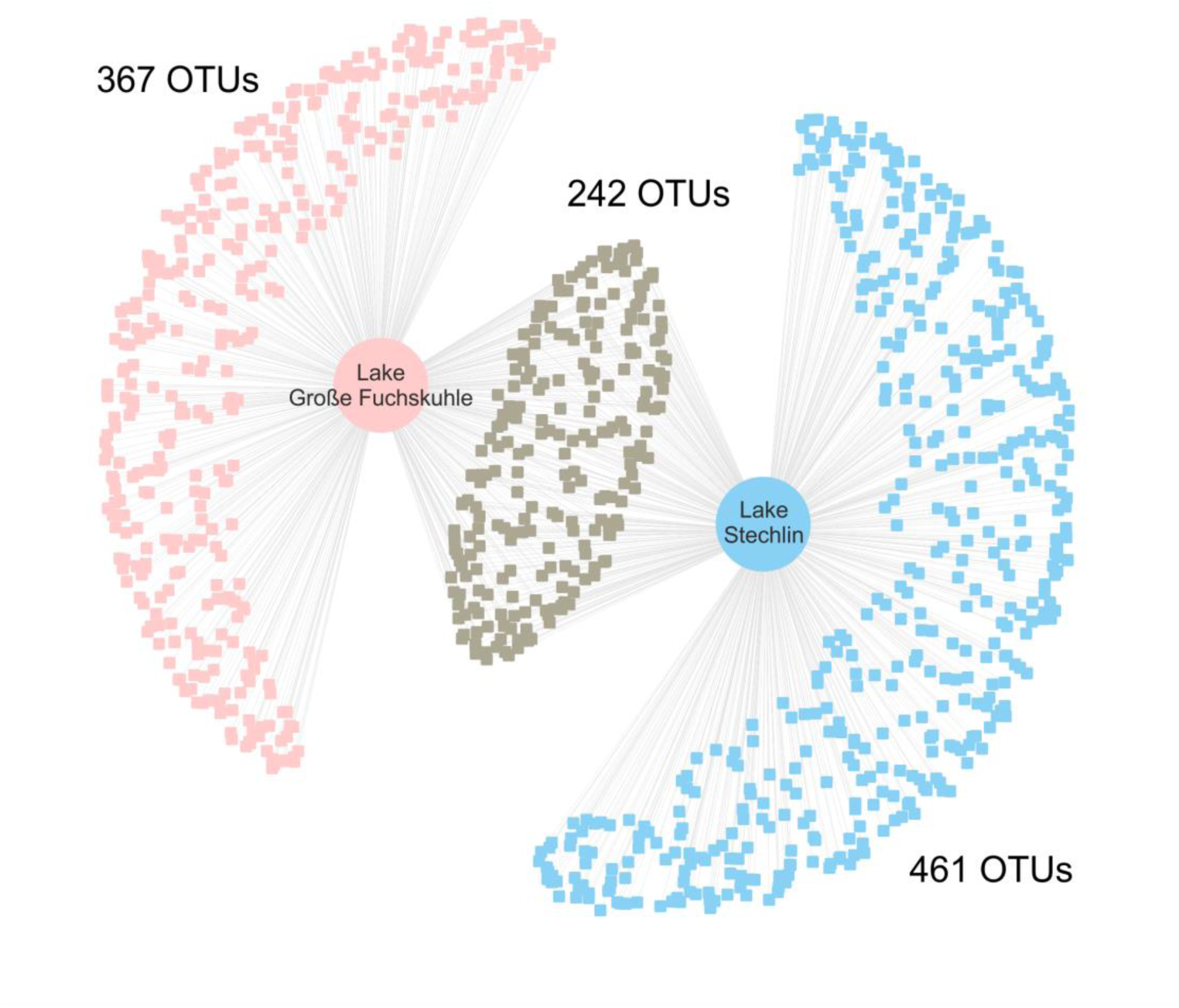
A network illustrating shared and unique fungal OTUs in the aquifers of the two sampling sites. Red colour nodes represent OTUs unique to groundwater boreholes close to Lake Grosse Fuchskuhle, while blue nodes represent OTUs unique to boreholes close to Lake Stechlin. Grey nodes indicate OTUs that are shared by both groundwater bodies.

Alpha diversity of fungal communities revealed a contrasting picture between the two groundwater sites. The species richness per sample ranged from 27 to 88 species, with significantly lower values in the near-shore shallow groundwater boreholes around Lake Stechlin compared to the deeper boreholes of the same lake or those around Lake Grosse Fuchskuhle (Fig. 6a, Table S2). Evenness revealed no significant difference between the groups and was generally low, with greater variation among the shallower boreholes, indicating that a few species are dominating the fungal community in the groundwater (Fig. 6b, Table S2).

**Fig 6:**
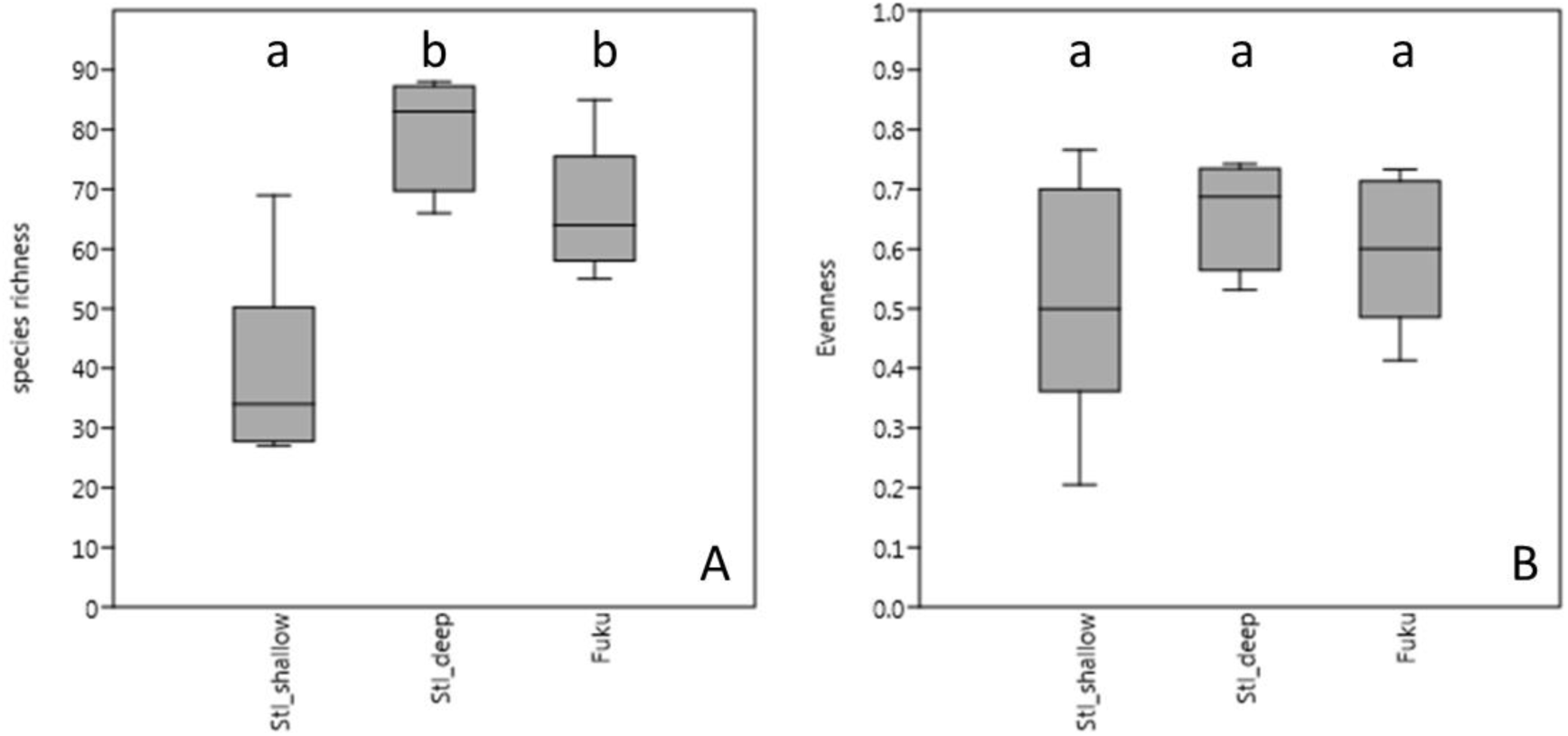
Species richness (a) and evenness (b) of fungal communities for the different sampling sites. Box plots show the median value (center line), the upper/lower quartiles (box edges) and maxima/minima (whiskers). Significance between the different groups is indicated by small letters above the boxes.

### Fungal isolates and their degradation capacity of complex, polymeric substances

Twenty seven fungal strains were isolated from three different boreholes covering a DOC concentration gradient (boreholes FX3, NW1, and NGW250). The isolates were highly diverse, and assigned to two fungal phyla (*Ascomycota* and *Zygomycota*), five classes (*Dothideomycetes, Eurotiomycetes, Leotiomycetes, Mucoromycotina, Sordariomycetes*), nine orders, and 14 different families. Figure 7 shows the phylogenetic relationship between the fungal isolates.

**Fig 7:**
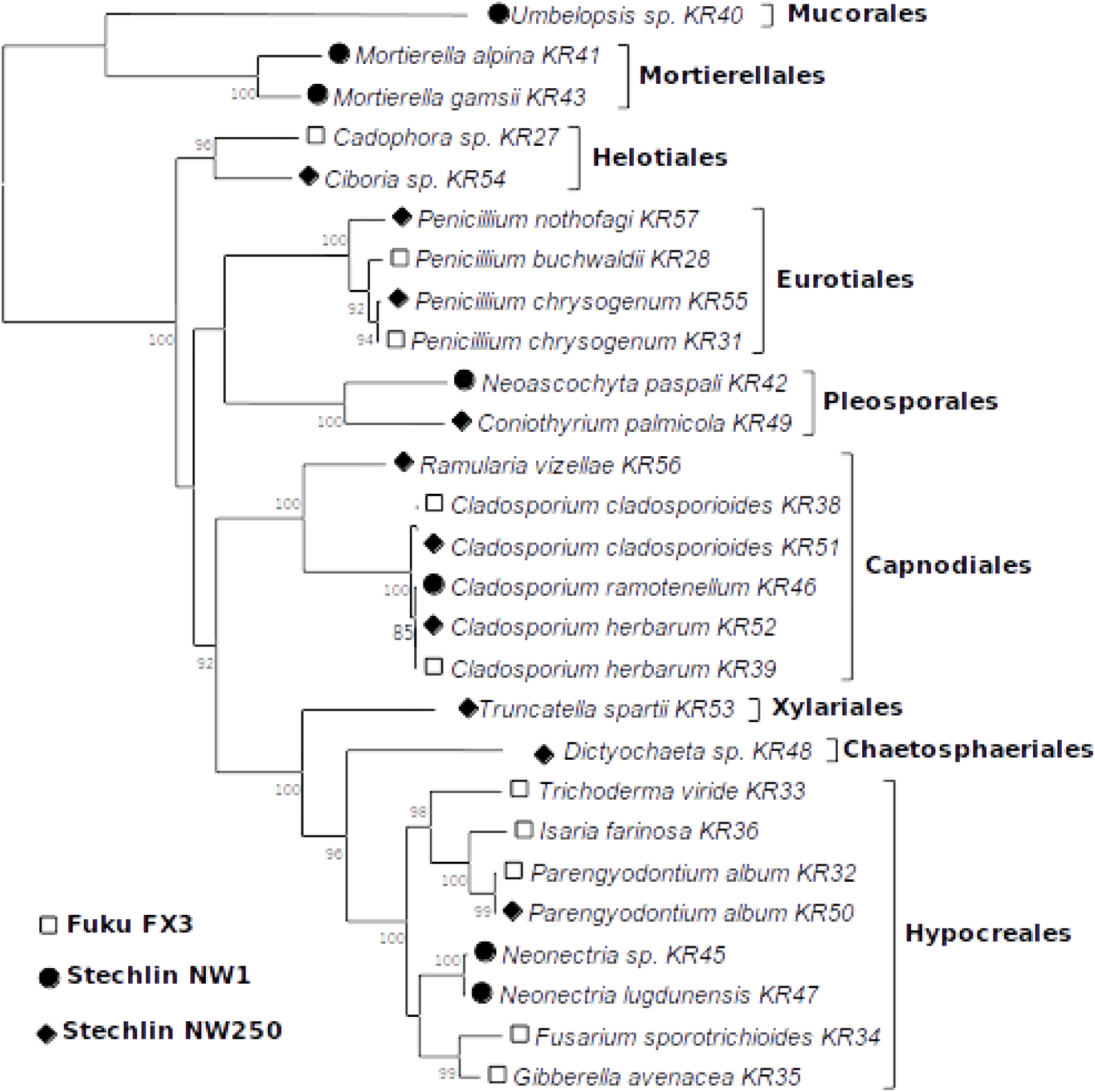
Maximum likelihood tree of 27 fungal isolates from three different groundwater sites based on a concatenated alignment of ITS1-5.8S-ITS2 and 28 rDNA gene regions (1195 bp). Bootstrap values higher than 70% are indicated on the nodes. Symbols represent the borehole origin, fungal orders are shown in bold.

The most abundant orders consisted of *Hypocreales* and *Capnodiales*, isolated from all three boreholes. All other orders were observed only in two of the sites: *Eurotiales* and *Heliotiales* in samples Fuku FX3 and Stechlin NW250, or *Pleosporales* in Stechlin NW1 and Stechlin NW250. Members of *Mucorales* and *Mortierellales* (both *Zygomycota*) were only isolated from Stechlin NW1. The most frequently isolated genus was *Cladosporium* which was isolated from all three boreholes. We were also able to find most of the isolated fungal strains in our Illumina sequencing data (see Table S3) and with most strains being found also in other boreholes investigated in this study.

Most of the isolates were able to degrade at least one of the polymeric substrates tested (Table 2). Three isolates showed a degradation capability for Bromocresol Green (Bromo), 12 isolates for Remazol Brilliant Blue (RBBR), 16 isolates for 2,2′-Azino-bis 3-ethylbenzothiazoline-6-sulfonic acid (ABTS) and 21 isolates for Congo Red (Congo), while 10 isolates showed activity on PolyR-478 (PolyR) and 12 on Toluidine (Tol). Most isolates could degrade at least one substrate, except strains KR40 (*Umbelopsis* sp.) and KR56 (*Ramularia* sp.) which did not show any capacity to degrade the tested substances. Three isolates from Lake Grosse Fuchskuhle FX3 (KR27 – *Cadophora* sp., KR35 – *Gibberella* sp., KR36 – *Isaria* sp.) were able to degrade all six tested substrates. We also observed that some isolates differed in their degradation capabilities despite having the same taxonomic classification (i.e. *Cladosporium herbarum* and *C. cladosporioides* isolated from boreholes Fuku FX3 and Stechlin NW250).

**Table 2:**
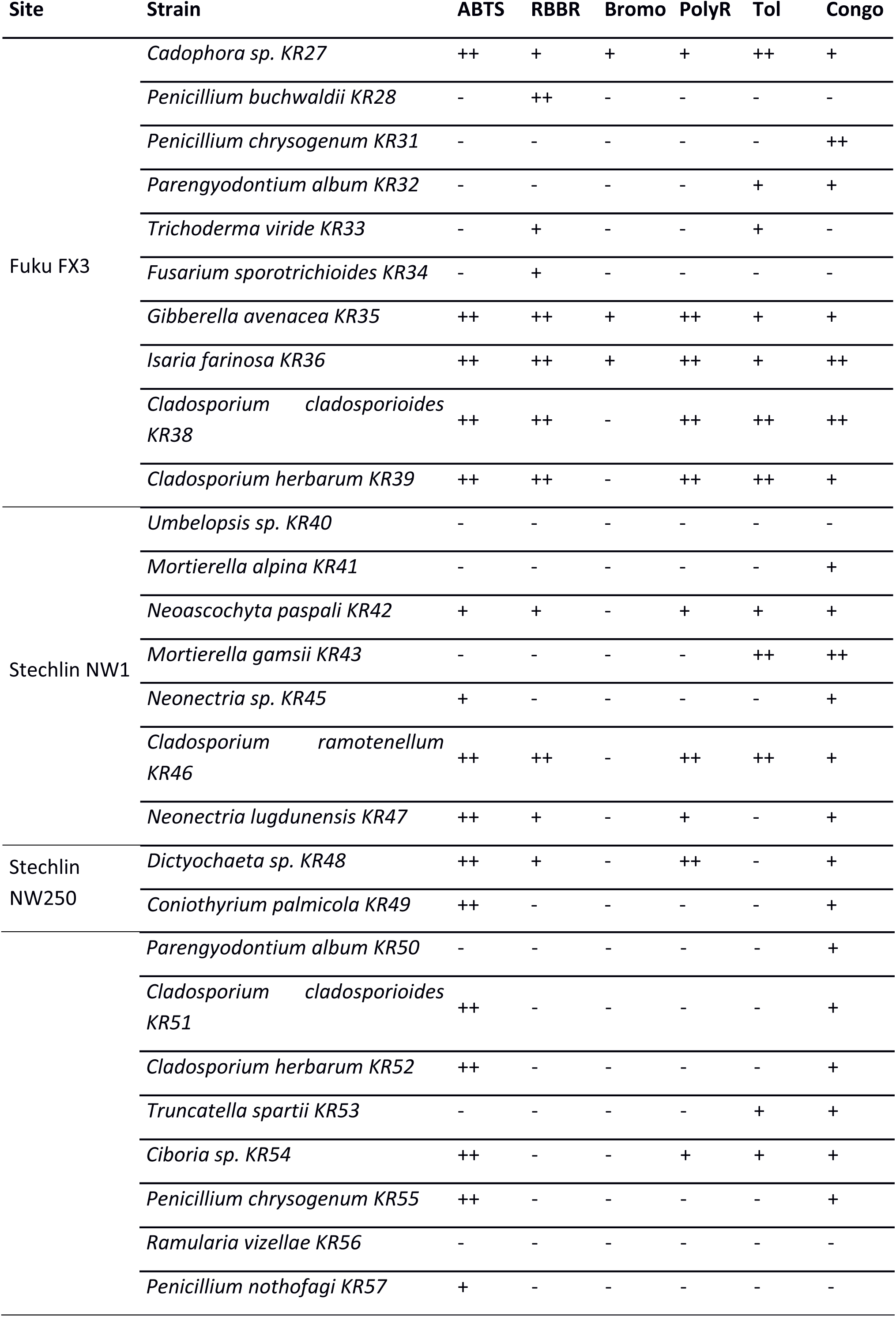
Results of the degradation assays for different polymeric substrates. Degradation positive activities are shown in a semi-quantitative scale: -, + and ++ represent no, low and high activities, respectively. Capacity of degrading 2,2′-Azino-bis 3-ethylbenzothiazoline-6-sulfonic acid (ABTS) was detected as a color change, demonstrating laccase activity. Decolorization of Remazol Brilliant Blue (RBBR) is related to lignin peroxidase activity, while decolorization of PolyR-478 (PolyR), Bromocresol Green (Bromo), Toluidine (Tol), and Congo Red (Congo) is related to degradation of polymeric, triarylmethane, and heterocyclic substrates, respectively.

## Discussion

Illumina amplicon sequencing and our isolation-based approach revealed a wide diversity of fungi in aquifers in the vicinity of two lakes with contrasting limnological features. We were able to demonstrate that groundwater can contain high concentrations of DOM, which varied in its optical properties such as SUVA_254_ indicating the occurrence of different chemical and microbiological transformation processes between collection sites. The majority of the groundwater fungal sequences were associated with *Ascomycota* and *Basidiomycota*, but also *Chytridiomycota, Cryptomycota* and *Zygomycota* were present. Most of the isolated fungal strains were able to degrade highly polymeric OM, confirming an active and important role of fungi in groundwater DOM degradation.

Although the role of fungi in terrestrial ecosystems is well recognized, their role in groundwaters has been little studied. Our results indicate that aquifer fungi have the same or similar ecological functions as in soil, i.e. the degradation of complex polymeric substances. Carbon quality and quantity in groundwater is greatly influenced by the sedimentary legacy of the surrounding ground and may explain differences in the biogeochemical parameters of the investigated boreholes (Artinger et al., 2000). The carbon content in aquifers has implications for fungal community assembly and associated functions, as fungi can degrade and modify a variety of organic carbon compounds including highly polymeric OM (Flaig, 1964; Dashtban et al., 2010; Harms et al., 2011). Fungi are also involved in various parts of the nitrogen cycle, e.g. by providing nitrogen to other members of the groundwater biota via “nitrogen mining” from organic macromolecules such as lignin (Hobbie et al., 2013). Under sub-and anoxic conditions, as often found in groundwater, they can use dissimilatory nitrate reduction pathways to gain energy (Takaya, 2002; Takasaki et al., 2004), possibly coupled with iron oxidation (Smith et al., 2017). These processes might be relevant because of the high ammonium and iron concentrations, and low dissolved oxygen concentrations, in several of the boreholes, particularly those close to Lake Grosse Fuchskuhle as well as the deepest borehole (250 m) close to Lake Stechlin.

All boreholes were characterized by the presence of *Ascomycota* and *Basidiomycota*. Members of these phyla are well known as degraders of dead complex organic matter in terrestrial and aquatic habitats (Baldrian et al., 2011; Duarte et al., 2015). Based on our results, including the diversity study and degradation assays, we suggest that these groundwater fungi might be involved in nutrient cycling as well as in carbon degradation and restructuring in aquifers (Rojas-Jiménez et al., 2017). However, some of the investigated boreholes showed a high proportion of *Chytridiomycota, Zygomycota* and/or *Cryptomycota* sequences. These fungal groups include saprotrophic, parasitic or even hyperparasitic members (White et al., 2006; Kagami et al., 2007; Gleason et al., 2014). Species of *Chytridiomycota* can infect a wide variety of phytoplankton species (e.g. diatoms, cyanobacteria, green algae; Kagami et al., 2007), but can also grow on dead organic matter, e.g. on pollen (Wurzbacher et al., 2014). Since active phytoplankton is most likely absent in groundwater, we propose two different lifestyles for the detected chytrids: 1. saprotrophes/phagotrophes using complex organic compounds for growth and metabolism and/or 2. parasites on groundwater meiofauna such as rotifers or nematodes (Deacon and Saxena, 1997; Glockling, 1998). The role of *Zygomycota* in groundwater is also largely unknown. Species of this phylum are known to have a saprophytic lifestyle, while others are parasites or can even actively catch rotifers and nematodes (Karling, 1936; Glockling, 1997). Indeed, very recently, it has been shown that they can control rotifer populations in activated sludge (Pajdak-Stós et al., 2016). Members of the *Zygomycota* might therefore play a similarly important role in regulating the groundwater fauna as suggested for other fungal groups (Nordbring-Hertz et al., 2011).

*Cryptomycota* are mainly represented by environmental sequences, and only a few isolates have been described to date (e.g. *Rozella allomyces*; Foust, 1937). The closest phylogenetic neighbours of the *Cryptomycota* sequences detected in our samples belong to sequences that were found in a pollen incubation experiment in Lake Stechlin (Wurzbacher et al., 2014) and seem to be related to the presence of *Chytridiomycota* and *Oomycota*. Both groups include well known saprophytes, i.e. as degraders of pollen. *Cryptomycota* are described as parasites of other fungi (Letcher et al., 2017), but can also infect *Oomycota* (Letcher et al., 2018). The (hyper)parasitic lifestyle suggests that they can have a strong influence on other trophic levels and might therefore play an important regulatory role for groundwater ecosystems.

Groundwaters and surface waters are often seen and managed as separate units, however, species exchange between surface and subsurface can occur (Brad et al., 2008). This is supported by the detection of a dominant fungal OTU (Otu0002) in our data set. This OTU is closely related to *Exobasidium* sp. (*Basidiomycota*) which is a known plant pathogen on *Vaccinium* sp. which is frequently growing in the sampling area. Since we used a DNA-based approach, we cannot fully exclude a potential transport of spores or mycelia through the soil substrate into the water-bearing formations. It has been shown in earlier studies that there is an interconnection of groundwater with surface waters, which can lead to the dispersion of fungi and other microorganisms even over large distances (Pyle et al., 1979; Perkins et al., 2015; Nawaz et al., 2016), although that is also dependent on the flow velocity, aquifer porosity and atmospheric pressure. Recently, Nawaz et al. (2018) used ITS mRNA to show that fungi are active and diverse in groundwater. They reported that the OTU overlap with an earlier DNA-based study (Nawaz et al., 2016) was only around 6%, which points to a difference between the overall and the active fungal community. However, this rather small overlap can be a consequence of comparing samples from different seasons and years, but also by precluding the detection of the non-and low-active as well as the low-abundant part of the fungal community when using a DNA-versus a RNA-based approach. Furthermore, differences in the used primer sets and the bioinformatic workflow make a direct comparison of the fungal datasets generated in the two studies difficult (Sinclair et al. 2015). Although we might have partly captured in our study of the non-active fungal fraction dwelling in groundwater by using a DNA-based sequencing approach, our study adds valuable information to the overall diversity, and by using an isolation approach it points to potential functions of fungi in aquifers (Hongsanan et al., 2018).

Despite the limitations of in vitro culturing methods, the majority of the obtained isolates were detected in our metabarcoding analysis (see Table S3), and more importantly, we confirmed that they can be active degraders of polymeric substances. The most frequently isolated genus was *Cladosporium*, while the most taxon-rich orders were *Hypocreales* and *Capnodiales*, which were isolated from all three boreholes. These two orders are known to contain several genera that are capable of degrading complex polymers, including cellulose and lignin (Rojas-Jiménez and Hernández, 2015), which was confirmed by our degradation assay. Further, our data indicates that differences in DOM composition between the two main groundwater sites were mainly due to variations in DOM aromaticity. On one hand, fungi can degrade humic substances and on the other hand they can actively increase the production of more aromatic compounds (Rojas-Jiménez et al., 2017). Although based on a reduced dataset, we noted that isolates from the groundwater close to Lake Grosse Fuchskuhle were able to degrade a much wider range of OM than those isolated from groundwater close to Lake Stechlin. Therefore, we suggest that fungi from Lake Grosse Fuchskuhle have a higher potential to increase DOM aromaticity via DOM transformation. Unfortunately, our dataset does not allow disentangling the transformation of humic acids from the synergy of processes occurring in the natural environment. Our data, however, provide evidence that groundwater systems with highly aromatic organic compounds can be well populated by fungi with a wide functional potential, in particular to degrade a variety of complex polymeric substances. As previously demonstrated, fungi are important players in the global carbon and nitrogen cycles (Krauss et al., 2011; Rütting et al., 2011); therefore, isolation and quantification of such processes in groundwater will improve our understanding of humic matter cycling also at the global scale.

Despite our limited knowledge about fungi in aquatic systems, and especially in groundwater, we revealed a highly diverse fungal community in different groundwater habitats in the vicinity of two lakes with contrasting limnological features. Our combined molecular and cultivation-based approaches demonstrate the potential role of aquatic fungi in groundwater habitats, especially for the cycling of organic matter by degrading complex organic compounds as well as in the ecological regulation of groundwater biota via parasitism. Some of the isolated fungi had the same taxonomic classification but differed in their degradation capabilities, pointing to functional variations in ecotypes or differences at strain level. In order to detect these functional differences also on the taxonomic level, the use of additional genetic markers is suggested for future investigations.

## Acknowledgements

We thank Christine Sturm and Jörg Lewandowski for providing sampling equipment and help during fieldwork as well as Uta Mallock for help with the carbon measurement, and Elke Mach for laboratory support. Luca Zoccarato is acknowledged for help with taxonomic classification and providing the Python script. The project was funded by the Leibniz Society SAW project MycoLink (SAW-2014-IGB).

## References

Allgaier M, Grossart H-P (2006) Seasonal dynamics and phylogenetic diversity of free-living and particle-associated bacterial communities in four lakes in northeastern Germany. Aquat Microb Ecol 45:115–128

Artinger R, Buckau G, Geyer S, Fritz P, Wolf M, Kim JI (2000) Characterization of groundwater humic substances: influence of sedimentary organic carbon. Appl Geochem 15:97–116

Baldrian P, Voříšková J, Dobiášová P, Merhautová V, Lisá L, Valášková V (2011) Production of extracellular enzymes and degradation of biopolymers by saprotrophic microfungi from the upper layers of forest soil. Plant Soil 338:111–125

Bertilsson S, Tranvik LJ (2000) Photochemical transformation of dissolved organic matter in lakes. Limnol Oceanogr 45:753–762

Brad T, Braster M, van Breukelen BM, van Straalen NM, Röling WF (2008) Eukaryotic diversity in an anaerobic aquifer polluted with landfill leachate. Appl Environ Microbiol 74:3959–3968

Burkert U, Ginzel G, Babenzien HD, Koschel R (2004) The hydrogeology of a catchment area and an artificially divided dystrophic lake – consequences for the limnology of Lake Fuchskuhle. Biogeochemistry 71:224–246

Cai W J, Wang Y, Krest J, Moore WS (2003) The geochemistry of dissolved inorganic carbon in a surficial groundwater aquifer in North Inlet, South Carolina, and the carbon fluxes to the coastal ocean. Geochim Cosmochim Ac 67:631–639

Casper SJ (1985) Lake Stechlin: A temperate oligotrophic lake. Dr W. Junk Publishers, Dordrecht

Chapelle F (2001) Ground-water microbiology and geochemistry. John Wiley & Sons

Chapelle FH, Bradley PM, Journey CA, McMahon PB (2013) Assessing the relative bioavailability of DOC in regional groundwater systems. Groundwater 51:363–372

Christmann D, Crouch S, Holland J, Timnick A (1980) Correction of right-angle molecular fluorescence measurements for absorption of fluorescence radiation. Anal Chem 52:291–295

Clipson N, Gleeson DB (2012) Fungal biogeochemistry: a central role in the environmental fate of lead. Curr Biol 22:R82–R84

Coble PG (1996) Characterization of marine and terrestrial DOM in seawater using excitation- mission matrix spectroscopy. Mar Chem 51:325–346

Crowther TW, Grossart H-P (2015) The role of bottom-up and top-down interactions in determining microbial and fungal diversity and function. In: Hanley TC, La Pierre KJ (eds) Trophic Ecology: Bottom-up and top-down interactions across aquatic and terrestrial systems. Cambridge University Press, pp 260–286

Csardi G, Nepusz T (2006) The igraph software package for complex network research. InterJournal, Complex Systems, 1695

Danielopol DL, Griebler C, Gunatilaka A, Notenboom J (2003) Present state and future prospects for groundwater ecosystems. Environ Conserv 30:104–130

Dashtban M, Schraft H, Syed TA, Qin W (2010) Fungal biodegradation and enzymatic modification of lignin. Int J Biochem Mol Biol 1:36–50

Deacon JW, Saxena G (1997) Orientated zoospore attachment and cyst germination in *Catenaria anguillulae*, a facultative endoparasite of nematodes. Mycol Res 101:513–522

Duarte S, Bärlocher F, Trabulo J, Cássio F, Pascoal C (2015) Stream-dwelling fungal decomposer communities along a gradient of eutrophication unraveled by 454 pyrosequencing. Fung Div 70:127–148

Flaig W (1964) Effects of micro-organisms in the transformation of lignin to humic substances. Geochim Cosmochim Ac 10-11:1523–1535

Foust FK (1937) A new species of *Rozella* parasitic on *Allomyces*. J Elisha Mitch Sci S 53:197–204

Gessner MO (1997) Fungal biomass, production and sporulation associated with particulate organic matter in streams. Limnetica 13:33–44

Ginzel G (1999) Hydrological investigations in the catchment area of Lake Stechlin and Lake Nehmitz. Berichte des IGB, Berlin, Heft 9, pp 43–60

Gleason FH, Lilje O, Marano AV, Sime-Ngando T, Sullivan BK, Kirchmair M, Neuhauser S (2014) Ecological functions of zoosporic hyperparasites. Front Microbiol 5:244

Glockling SL (1997) *Zoophagus cornus*: a new species from Japan. Mycol Res 101:1179–1182

Glockling SL (1998) Isolation of a new species of rotifer-attacking *Olpidium*. Mycol Res 102:206–208

Graham PW, Baker A, Andersen MS (2015) Dissolved organic carbon mobilisation in a groundwater system stressed by pumping. Sci Rep 5:18487

Grossart H-P, Simon M (2007) Interactions of planktonic algae and bacteria: effects on algal growth and organic matter dynamics. Aquat Microb Ecol 47:163–176

Grossart H-P, Wurzbacher C, James TY, Kagami M (2016) Discovery of dark matter fungi in aquatic ecosystems demands a reappraisal of the phylogeny and ecology of zoosporic fungi. Fung Ecol 19:28–38

Grossart H-P, Rojas-Jiménez K (2016) Aquatic fungi: targeting the forgotten in microbial ecology. Curr Opin Microbiol 31:140–145

Hall T (1999) Bioedit: a user-friendly biological sequence alignment editor and analysis program for windows 95/98/NT. Nucleic Acids Symp Ser 41:95–98

Harms H, Schlosser D, Wick LY (2011) Untapped potential: exploiting fungi in bioremediation of hazardous chemicals. Nat Rev Microbiol 9:177 – 192

Helms JR, Stubbins A, Ritchie JD, Minor EC, Kieber DJ, Mopper K (2008) Absorption spectral slopes and slope ratios as indicators of molecular weight, source, and photobleaching of chromophoric dissolved organic matter. Limnol Oceanogr 53:955–969

Hobbie EA, Ouimette AP, Schuur EAG, Kierstead D, Trappe JM, Bendiksen K, Ohenoja E (2013) Radiocarbon evidence for the mining of organic nitrogen from soil by mycorrhizal fungi. Biogeochemistry 114:381–389

Holzbecher E, Nützmann G, Ginzel G (1999) Water and component mass balances in the catchment of Lake Stechlin. In: Leibundgut C, McDonnell J, Schultz G (ed) Integrated methods in catchment hydrology: tracer, remote sensing and new hydrometric techniques. IAHS Press, Wallingford, pp 37–44

Hongsanan S, Jeewon R, Purahong W, Xie N, Liu J-K, Jayawardena RS, Ekanayaka AH, Dissanayake A, Raspé O, Hyde KD, Stadler M, Peršoh D (2018) Can we use environmental DNA as holotypes? Fung Div 92:1–30

Hutalle-Schmelzer KML, Zwirnmann E, Krüger A, Grossart H-P (2010) Enrichment and cultivation of pelagic bacteria from a humic lake using phenol and humic matter additions. FEMS Microbiol Ecol 72:58–73

Jardine PM, McCarthy JF, Weber NL (1989) Mechanisms of dissolved organic carbon adsorption on soil. Soil Sci Soc Am J 53:1378–1385

Jørgensen NOG, Stepanauskas R (2009) Biomass of pelagic fungi in Baltic rivers. Hydrobiologia 623:105–112

Kaboth U, Rechlin B, Ginzel G (2008) Besteht für unsere Seen eine geogene Versalzungsgefahr? Hydrochemisch-genetische Untersuchungen von Speisungsbedingungen an Seen im Naturpark Stechlin. Brandenburgische Geowissenschaftliche Beiträge 15:69–79

Kagami M, de Bruin A, Ibelings BW, Van Donk E (2007) Parasitic chytrids: their effects on phytoplankton communities and food-web dynamics. Hydrobiologia 578:113–129

Karling JS (1936) A new predaceous fungus. Mycologia 28:307–320

Katoh K, Standley DM (2013) MAFFT multiple sequence alignment software version 7: improvements in performance and usability. Mol Biol Evol 30:772–780

Korbel K, Chariton A, Stephenson S, Greenfield P, Hose GC (2017) Wells provide a distorted view of life in the aquifer: implications for sampling, monitoring and assessment of groundwater ecosystems. Sci Rep 7:40702

Krauss GJ, Solé M, Krauss G, Schlosser D, Wesenberg D, Bärlocher F (2011) Fungi in freshwaters: ecology, physiology and biochemical potential. FEMS Microbiol Rev 35:620–651

Kumar S, Stecher G, Tamura K (2016) MEGA7: Molecular Evolutionary Genetics Analysis version 7.0 for bigger datasets. Mol Biol Evol 33:1870–1874

Lategan MJ, Hose GC (2014) Development of a groundwater fungal strain as a tool for toxicity assessment. Environ Toxicol Chem 33:2826–2834

Lategan MJ, Torpy F, Newby S, Stephenson S, Hose GC (2012) Fungal diversity of shallow aquifers in southeastern Australia. Geomicrobiol J 29:352–361

Letcher PM, Longcore JE, Quandt CA, da Silva Leite D, James TY, Powell MJ (2017) Morphological, molecular and ultrastructural characterization of *Rozella rhizoclosmatii*, a new species in *Cryptomycota*. Fung Ecol 121:1–10

Letcher PM, Longcore JE, James TY, Leite DS, Simmons DR & Powell MJ (2018) Morphology, Ultrastructure, and molecular phylogeny of *Rozella multimorpha*, a new species in *Cryptomycota*. J Eukaryot Microbiol 65:180–190

Li J, Wang Y, Guo W, Xie X, Zhang L, Liu Y, Kong S (2014) Iodine mobilization in groundwater system at Datong basin, China: evidence from hydrochemistry and fluorescence characteristics. Sci Total Environ 468–469:738–745

Liedtke H (1975) Die nordischen Vereisungen in Mitteleuropa. Erläuterungen zu einer farbigen Übersichtskarte im Maßstab 1: 1000000. Bundesforschungsanstalt für Landeskunde und Raumordnung

Lochmüller CH, Saavedra SS (1986) Conformational changes in a soil fulvic acid measured by time-dependent fluorescence depolarization. Anal Chem 58:1978–1981

Macpherson GL (2009) CO2 distribution in groundwater and the impact of groundwater extraction on the global C cycle. Chem Geol 264:328–336

Maher DT, Santos IR, Golsby-Smith L, Gleeson J, Eyre BD (2013) Groundwater-derived dissolved inorganic and organic carbon exports from a mangrove tidal creek: the missing mangrove carbon sink? Limnol Oceanogr 58:475–488

Mitch WJ, Gosselink JG (2007) Wetland biogeochemistry. In: Mitch WJ, Gosselink JG (eds) Wetlands, 4th edn. Wiley, Hoboken, New Jersey, pp 163–206

Mladenov N, Zheng Y, Miller MP, Nemergut DR, Legg T, Simone B, Hageman C, Rahman MM, Ahmed KM, Mcknight DM (2010) Dissolved organic matter sources and consequences for iron and arsenic mobilization in Bangladesh aquifers. Environ Sci Technol 44:123–128

Mueller RC, Gallegos-Graves L, Kuske CR (2016) A new fungal large subunit ribosomal RNA primer for high-throughput sequencing surveys. FEMS Microbiol Ecol 92:fiv153

Murphy KR, Stedmon CA, Graeber D, Bro R (2013) Fluorescence spectroscopy and multi-way techniques. PARAFAC. Anal Methods 5:6557–6566

Nawaz A, Purahong W, Lehmann R, Herrmann M, Küsel K, Totsche KU, Buscot F, Wubet T (2016) Superimposed pristine limestone aquifers with marked hydrochemical differences exhibit distinct fungal communities. Front Microbiol 7:666

Nawaz A, Purahong W, Lehmann R, Herrmann M, Totsche KU, Küsel K, Wubet T, Buscot F (2018) First insights into the living groundwater mycobiome of the terrestrial biogeosphere. Water Res 145:50–61

Nercessian O, Noyes E, Kalyuzhnaya MG, Lidstrom ME, Chistoserdova L (2005) Bacterial populations active in metabolism of C1 compounds in the sediment of Lake Washington, a freshwater lake. Appl Environ Microbiol 71:6885–6899

Nordbring-Hertz B, Jansson H & Tunlid A (2011) Nematophagous Fungi. eLS, https://doi.org/10.1002/9780470015902.a0000374.pub3.

Ohno T (2002) Fluorescence inner-filtering correction for determining the humification index of dissolved organic matter. Environ Sci Technol 36:742–746

Oksanen J, Blanchet F, Kindt R, Legendre P, Minchin P, O’Hara R, Simpson G, Solymos P, Henry M, Stevens H (2017) Vegan: Community Ecology Package. R-package version 2.4-0

Pajdak-Stós A, Wazny R, Fiałkowska E (2016) Can a predatory fungus (*Zoophagus* sp.) endanger the rotifer populations in activated sludge? Fung Ecol 23:75–78

Pearson WR (2002) Finding protein and nucleotide similarities with FASTA. Current Protocols in Bioinformatics 53:3.9.1–3.9.25

Perkins AK, Santos IR, Sadat-Noori M, Gatland JR, Maher DT (2015) Groundwater seepage as a driver of CO2 evasion in a coastal lake (Lake Ainsworth, NSW, Australia). Environ Earth Sci 74:779– 792

Peter S, Shen Y, Kaiser K, Benner R, Durisch-Kaiser E (2012) Bioavailability and diagenetic state of dissolved organic matter in riparian groundwater. J Geophys Res-Biogeo 117:G04006

Price MN, Dehal PS, Arkin AP (2010) FastTree 2–approximately maximum-likelihood trees for large alignments. PLoS One 5:e9490

Pruesse E, Peplies J, Glöckner FO (2012) SINA: accurate high-throughput multiple sequence alignment of ribosomal RNA genes. Bioinformatics 28:1823–1829

Pyle BH, Sinton LW, Noonan MJ & McNabb JF (1979) The movement of micro-organisms in groundwater. In: Technical Group on Water: Proceedings of a Symposium Held by the Group in Conjunction with the Annual Conference of the N.Z.I.E., Wellington, 5:465–481

R-Core-Team (2017) R: A language and environment for statistical computing. R Foundation for Statistical Computing, Vienna, Austria

Rochelle-Newall EJ, Fisher TR (2002) Chromophoric dissolved organic matter and dissolved organic carbon in Chesapeake Bay. Mar Chem 77:23–41

Rojas-Jiménez K & Hernández M (2015) Isolation of fungi and bacteria associated with the guts of tropical wood-feeding coleoptera and determination of their lignocellulolytic activities. Int J Microbiol 2015:285018

Rojas-Jiménez K, Fonvielle JA, Ma H, Grossart H-P (2017) Transformation of humic substances by the freshwater Ascomycete *Cladosporium* sp.. Limnol Oceanogr 62:1955–1962

Rütting T, Boeckx P, Müller C, Klemedtsson L (2011) Assessment of the importance of dissimilatory nitrate reduction to ammonium for the terrestrial nitrogen cycle. Biogeosciences 8:1779–1791

Santos IR, Maher DT, Eyre BD (2012) Coupling automated radon and carbon dioxide measurements in coastal waters. Environ Sci Technol 46:7685–7691

Shannon P, Markiel A, Ozier O, Baliga NS, Wang JT, Ramage D, Amin N, Schwikowski B, Ideker T (2003) Cytoscape: a software environment for integrated models of biomolecular interaction networks. Genome Res 13:2498–2504

Shen Y, Chapelle FH, Strom EW, Benner R (2015) Origins and bioavailability of dissolved organic matter in groundwater. Biogeochemistry 122:61–78

Sinclair L, Osman OA, Bertilsson S, Eiler A (2015) Microbial community composition and diversity via 16S rRNA gene amplicons: evaluating the Illumina platform. PLoS ONE 10:e0116955

Smith RL, Kent DB, Repert DA & Böhlke JK (2017) Anoxic nitrate reduction coupled with iron oxidation and attenuation of dissolved arsenic and phosphate in a sand and gravel aquifer. Geochim Cosmochim Ac 196:102–120

Sohlberg E, Bomberg M, Miettinen H, Nyyssönen M, Salavirta H, Vikman M, Itävaara M (2015) Revealing the unexplored fungal communities in deep groundwater of crystalline bedrock fracture zones in Olkiluoto, Finland. Front Microbiol 6:573

Stedmon CA, Markager S, Bro R (2003) Tracing dissolved organic matter in aquatic environments using a new approach to fluorescence spectroscopy. Mar Chem 82:239–254

Takaya N (2002) Dissimilatory nitrate reduction metabolisms and their control in fungi. J Biosci Bioeng 94:506–510

Takasaki K, Shoun H, Yamaguchi M, Takeo K, Nakamura A, Hoshino T, Takaya N (2004) Fungal ammonia fermentation, a novel metabolic mechanism that couples the dissimilatory and assimilatory pathways of both nitrate and ethanol - role of acetyl CoA synthetase in anaerobic ATP synthesis. J Biol Chem 279:12414–12420

Treseder KK, Lennon JT (2015) Fungal traits that drive ecosystem dynamics on land. Microbiol Mol Biol Rev 79:243–262

Vilgalys R, Hester M (1990) Rapid genetic identification and mapping of enzymatically amplified ribosomal DNA from several Cryptococcus species J Bacteriol 172:4238–4246

Wang Q, Garrity GM, Tiedje JM, Cole JR (2007) Naïve bayesian classifier for rapid assignment of rRNA sequences into the new bacterial taxonomy. Appl Environ Microbiol 73:5261–5267

Weishaar JL, Aiken GR, Bergamaschi BA, Fram MS, Fugii R, Mopper K (2003) Evaluation of specific ultraviolet absorbance as an indicator of the chemical composition and reactivity of dissolved organic carbon. Environ Sci Technol 37:4702–4708

White TJ, Bruns TD, Lee S, Taylor J (1990) Amplification and direct sequencing of fungal ribosomal RNA genes for phylogenetics. In: Innis MA, Gelfand DH, Sninsky JJ, White TJ (eds) PCR protocols: a guide to methods and applications. Academic Press, New York, pp 315–322

White MM, James TY, O’Donnell K, Cafaro MJ, Tanabe Y, Sugiyama J (2006) Phylogeny of the *Zygomycota* based on nuclear ribosomal sequence data. Mycologia 98:872–884

Williams CJ, Yamashita Y, Wilson HF, Jaffé R, Xenopoulos MA (2010) Unraveling the role of land use and microbial activity in shaping dissolved organic matter characteristics in stream ecosystems. Limnol Oceanogr 55:1159–1171

Wilson HF, Xenopoulos MA (2009) Effects of agricultural land use on the composition of fluvial dissolved organic matter. Nat Geosci 2:37–41

Wurzbacher CM, Bärlocher F, Grossart H-P (2010) Fungi in lake ecosystems. Aquat Microb Ecol 59:125–149

Wurzbacher C, Rösel S, Rychła A, Grossart H-P (2014) Importance of saprotrophic freshwater fungi for pollen degradation. PLoS One 9:e94643

Zhang Y, Qin B, Zhu G, Zhang L, Yang L (2007) Chromophoric dissolved organic matter (CDOM) absorption characteristics in relation to fluorescence in Lake Taihu, China, a large shallow subtropical lake. Hydrobiologia 581:43–52

