## Supplementary Information for "Highly diverse fungal communities in carbon-rich aquifers of two contrasting lakes in Northeast Germany"

**Journal: Fungal Ecology**

Anita Perkins^1,2^*, Lars Ganzert^3,4,5^*^#^, Keilor Rojas-Jiménez^3,6^, Jeremy Fonvielle^3^, Grant C. Hose^1^, Hans-Peter Grossart^3,7#^

*these authors contributed equally to the manuscript

^1^Department of Biological Sciences, Macquarie University, New South Wales, Australia

^2^Present address: Centre for Coastal Biogeochemistry, School of Environmental Science and Engineering, Southern Cross University, 2480 Lismore, New South Wales, Australia

^3^Leibniz Institute for Freshwater Ecology and Inland Fisheries (IGB), Experimental Limnology, 16775 Neuglobsow, Germany

^4^GFZ German Research Centre for Geosciences, Helmholtz Centre Potsdam, Section 5.3 Geomicrobiology, Telegrafenberg C-422, 14473 Potsdam, Germany

^5^Experimental Phycology and Culture Collection of Algae (SAG), University of Göttingen, Nikolausberger Weg 18, 37073 Göttingen, Germany

^6^Escuela de Biologia, Universidad de Costa Rica, 11501 San Jose, Costa Rica

^7^University of Potsdam, Institute of Biochemistry and Biology, Maulbeerallee 2, 14469 Potsdam, Germany

Supplementary Information (SI)

Material and Methods

*Study Site*

The main geological formations marking the landscapes are end moraine, ground moraine, outwash, glacial spillway, eskers, kames and drumlins, channels and lakes (Liedtke 1975).

Borehole NGW250 (located at Lake Stechlin) is 250 m deep and accesses a layer of ancient marine sediments that contain coal and mud (Kaboth et al. 2008) (see Fig. S1). The above rupelton clay layer can reach 80 m thickness in some places and is of major hydrogeological significance, acting as a barrier between the ancient sea layer beneath and the freshwater reservoir above (Kaboth et al. 2008). Salinity in the ancient sea layer is up to 200 g L^-1^, much higher than the surrounding shallower wells NGW32, NGW80 and NGW160 (Kaboth et al. 2008).

Boreholes were drilled by Baugrund Berlin Bohrgesellschaft mbH in 1997 and by the Landesamt für Bergbau, Geologie und Rohstoffe (LBGR) in 2010. All boreholes were constructed of PVC pipes, except boreholes NGW which are made of stainless steel pipes. The diameter of the bores ranged from 6 to 15 cm.

*Optical characterisation of dissolved organic matter (DOM)*

DOM aromaticity (SUVA_254_) was obtained from the specific UV absorbance at 254 nm divided by the DOC concentration (Weishaar et al. 2003). We determined the ratio (hereafter noted S_r_) between the slope of decadal absorbance at short wavelength (from 275 to 295 nm) and the slope of decadal absorbance at long wavelength (from 350 to 400 nm) to appraise DOM molecular weight (Helms et al. 2008). We computed the ratio between decadal absorbance at 250 to 365 nm (E_2_:E_3_) to corroborate S_r_. The humification index (HIX) was computed by dividing the area of emission from 435 to 480 nm with the area of emission from 300 to 445 nm, at a constant excitation of 254 nm (Ohno 2002). The freshness of DOM (ß/α) was estimated as the maximal fluorescence intensity for an emission at 380 nm and an excitation at 310 nm (ß) and the maximal fluorescence intensity between emission at 420 and 435 nm (α) for a fixed excitation at 310 nm (Wilson et al. 2009). Individual peaks representing the signature of humic matter and protein-like DOM were retrieved as previously described (Lochmüller and Saavedra 1986; Coble 1996; Stedmon et al. 2003).

*DNA extraction*

Briefly, filters were homogenized in a bead beater using glass/zirconia beads (0.1, 0.7 and 3 mm), plus 600 µL extraction buffer (10% CTAB in 1.6 M NaCl mixed 1:1 with 240 mM K_2_HPO_4_/KH_2_PO_4_ buffer), 600 µL phenol:chloroform:isoamylalcohol (25:24:1) and 60 µL each 10% sodium dodecyl sulfate (SDS) and 10% N-lauroyl sarcosine. The aqueous phase was mixed with an equal volume of chloroform-isoamylalcohol (24:1) to remove any residual phenol. DNA was precipitated with 30% PEG 6000 in 1.6 M NaCl and the addition of 1 µL LPA (Sigma), followed by a centrifugation step. The resulting pellet was washed with 1 mL ice-cold ethanol (70%), dried briefly at 37°C, re-suspended in 50 µL PCR water and stored at -80°C until further processing.

Figure S1: Example of the color of the different groundwater samples. Samples from around Lake Grosse Fuchskuhle were yellow-brownish (left side of the picture) while samples from boreholes around Lake Stechlin were mostly colorless or only slightly yellowish (middle and right side). An exception is the borehole NGW250 which had a charcoal-like color (in front).

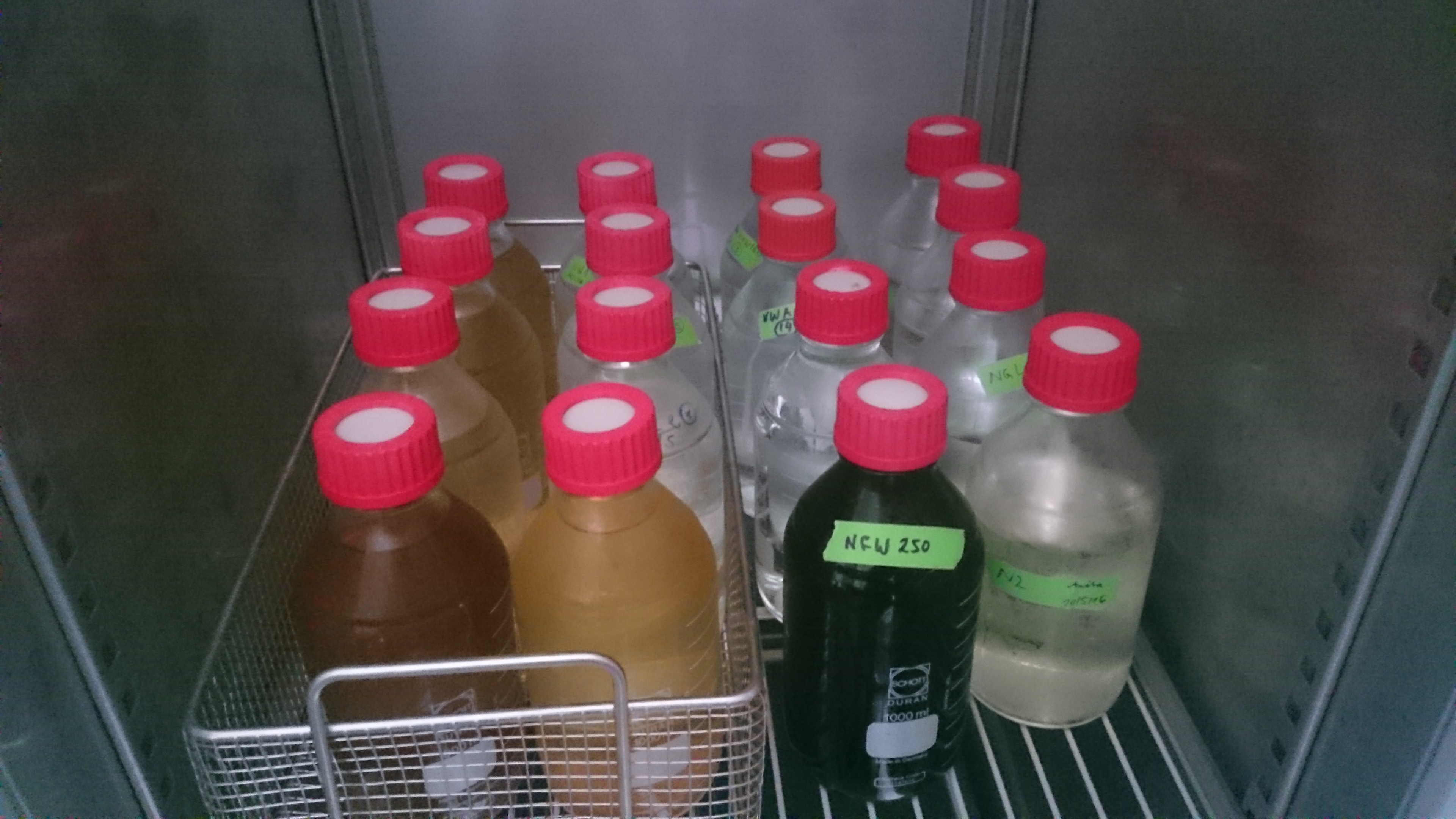

Table S3: Phylogenetic comparison between the fungal isolates and their closest environmental OTU match based on high-throughput sequencing reads from this study (incl. sequence similarity).

| **Isolate** | **closest OTU match** | **Similarity** |
| --- | --- | --- |
| *Cadophora* sp. KR27 | Otu1328 | 99.3% |
| *Penicillium buchwaldii* KR28 | Otu0120 | 97.4% |
| *Penicillium chrysogenum* KR31 | Otu0120 | 100% |
| *Parengyodontium album* KR32 | Otu0004 | 99.6% |
| *Trichoderma viride* KR33 | Otu0787 | 99.3% |
| *Fusarium sporotrichioides* KR34 | Otu0151 | 97.4% |
| *Gibberella avenacea* KR35 | Otu0151 | 100% |
| *Isaria farinosa* KR36 | Otu0075 | 99.6% |
| *Cladosporium cladosporioides* KR38 | Otu0019 | 100% |
| *Cladosporium herbarum* KR39 | Otu0019 | 100% |
| *Umbelopsis* sp. KR40 | Otu2343 | 92.8% |
| *Mortierella alpina* KR41 | Otu0246 | 100% |
| *Neoascochyta paspali* KR42 | Otu1529 | 95.2% |
| *Mortierella gamsii* KR43 | Otu1858 | 98.8% |
| *Neonectria* sp. KR45 | Otu0254 | 100% |
| *Cladosporium ramotenellum* KR46 | Otu0019 | 100% |
| *Neonectria lugdunensis* KR47 | Otu0254 | 100% |
| *Dictyochaeta* sp. KR48 | Otu0720 | 100% |
| *Coniothyrium palmicola* KR49 | Otu0105 | 100% |
| *Parengyodontium album* KR50 | Otu0004 | 100% |
| *Cladosporium cladosporioides* KR51 | Otu0019 | 100% |
| *Cladosporium herbarum* KR52 | Otu0019 | 100% |
| *Truncatella spartii* KR53 | Otu0077 | 100% |
| *Ciboria* sp. KR54 | Otu0372 | 94.5% |
| *Penicillium chrysogenum* KR55 | Otu0120 | 99.6% |
| *Ramularia vizellae* KR56 | Otu0010 | 100% |
| *Penicillium nothofagi* KR57 | Otu0243 | 100% |
